# Deletion of arrestin-3 does not improve compulsive drug-seeking behavior in a longitudinal paradigm of oral morphine self-administration

**DOI:** 10.1101/2023.03.30.534994

**Authors:** Sarah Warren Gooding, Lindsey Felth, Randi Foxall, Zachary Rosa, Kyle Ireton, Izabella Sall, Joshua Gipoor, Anirudh Gaur, Madeline King, Noah Dirks, Cheryl A. Whistler, Jennifer L. Whistler

## Abstract

Opioid drugs are potent analgesics that mimic the endogenous opioid peptides, endorphins and enkephalins, by activating the µ-opioid receptor. Opioid use is limited by side effects, including significant risk of opioid use disorder. Improvement of the effect/side effect profile of opioid medications is a key pursuit of opioid research, yet there is no consensus on how to achieve this goal. One hypothesis is that the degree of arrestin-3 recruitment to the µ-opioid receptor impacts therapeutic utility. However, it is not clear whether increased or decreased interaction of the µ-opioid receptor with arrestin-3 would reduce compulsive drug-seeking. To examine this question, we utilized three genotypes of mice with varying abilities to recruit arrestin-3 to the µ-opioid receptor in response to morphine in a novel longitudinal operant self-administration model. We demonstrate that arrestin-3 knockout and wild type mice have highly variable drug-seeking behavior with few genotype differences. In contrast, in mice where the µ-opioid receptor strongly recruits arrestin-3, drug-seeking behavior is much less varied. We created a quantitative method to define compulsivity in drug-seeking and found that mice lacking arrestin-3 were more likely to meet the criteria for compulsivity whereas mice with enhanced arrestin-3 recruitment did not develop a compulsive phenotype. Our data suggest that opioids that engage both G protein and arrestin-3, recapitulating the endogenous signaling pattern, will reduce abuse liability.

## INTRODUCTION

Opioids are powerful analgesic drugs that remain essential for the treatment of severe pain. Despite their therapeutic utility, opioid use can precipitate opioid use disorder (OUD). While most individuals who take opioids do not develop an OUD, over 2% of Americans age 12 and older meet the OUD diagnostic criteria (1) driving a major public health crisis, particularly with accidental overdose. Despite significant research efforts and billions of dollars invested, the development of an opioid with reduced abuse liability has been ultimately unsuccessful (2).This lack of success can be attributed in part to an incomplete understanding of how opioid signaling contributes to the physiological and behavioral components of OUD.

Opioid analgesia is primarily mediated by activation of the µ-opioid receptor (MOR), a G protein-coupled receptor (GPCR) (3). Endogenous opioid peptides, endorphins and enkephalins, bind and activate MOR to promote signaling to the G_i/o/z_ G protein effectors. G protein signaling from these peptide-occupied MORs is then titrated by a cascade of events that includes phosphorylation of the MOR by GPCR kinases (GRKs) (4, 5) and recruitment of the arrestin-3 (β-arrestin-2) effector to the phosphorylated receptor (6). Arrestin-3 recruitment not only uncouples MOR from its G protein but also promotes MOR endocytosis (7, 8). Endocytosed MORs are then dephosphorylated and recycled to the plasma membrane where they can bind ligand and initiate another cycle of signal transduction (9, 10). Activation of the MOR by opioid drugs, including morphine and all its derivatives, promotes G protein signaling like endogenous ligands. However, morphine-activated receptors only weakly engage the GRK and arrestin-3 effectors (4, 11, 12). This is because the morphine-activated MOR is phosphorylated on only one of the four residues (5) that are phosphorylated when the receptor is activated by an endogenous opioid. To denote this difference in MOR signaling by peptide or morphine occupied receptors, we refer to endogenous opioid peptides as balanced ligands: those that potently engage both the G protein and arrestin effectors. Small molecule opioid drugs are more biased: they more strongly engage G protein signaling in many cell types.

The impacts of biased and balanced signaling on the effect/side effect profile of opioid analgesics has been interrogated since the original discovery that morphine does not promote significant MOR endocytosis (13, 14). Decades later, there remains little consensus on the role of arrestin-3 recruitment in opioid side effects because both eliminating arrestin-3 recruitment and enhancing arrestin-3 recruitment reduces some of the side effects of morphine and strengthens its analgesic effects. Mice without the arrestin-3 gene (Arr3-KO) were reported to show increased analgesia (15), reduced tolerance (16), and reduced respiratory depression and constipation (17) in response to morphine compared to wild type (WT) mice. Likewise, knock-in mice where the MOR is replaced by a mutant receptor which cannot be phosphorylated by GRKs (MOR 11S/T-A) are also reported to show improved analgesia and reduced analgesic tolerance but no difference in respiratory depression in response to morphine (18). These data would suggest that removing arrestin-3 engagement improves analgesic utility. However, mice with a chimeric MOR that is an improved substrate for GRKs and have enhanced arrestin-3 recruitment (RMOR mice, for recycling MOR) also show enhanced analgesia (19) and reduced analgesic tolerance to morphine with no change in respiratory depression (20). In conditioned place preference (CPP) paradigms, both decreasing (Arr3-KO mice) (21) and increasing (RMOR knock-in mice) (22) arrestin-3 recruitment increases the potency of morphine reward. Finally, dependence, defined as physical and/or affective signs of distress upon the removal of drug, is another negative side effect of opioid use and a key component of OUDs. Both mouse lines deficient in arrestin-3 recruitment (Arr3-KO, MOR 11S/T-A) show intact or exacerbated morphine withdrawal signs, indicating that they still develop dependence (16, 18). In contrast, RMOR mice show neither physical (19) nor affective (22) signs of dependence upon withdrawal from morphine. This battery of conflicting results has left the field divided on the best therapeutic strategy for new opioid drugs.

In humans, OUD is a syndrome defined by a constellation of phenotypes that include loss of control in drug-seeking behavior, craving, and relapse as well as physiological tolerance and dependence. We have previously reported a three-phase operant self-administration paradigm that models aspects of compulsive drug-seeking in mice: escalation of drug-seeking (loss of control), failure to extinguish drug-seeking (craving), and reinstatement after prolonged abstinence (relapse). Using this model, we demonstrate that some WT but no RMOR mice become compulsive drug-seekers with time (22). However, it is not known how eliminating arrestin-3 impacts compulsive drug-seeking behavior. Since many of the side effects of opioids are improved with both the enhancement and the elimination of MOR-arrestin-3 interaction, its impact on drug-seeking is difficult to predict. We utilized a version of our compulsive drug-seeking model in three genotypes: WT, Arr3-KO, and RMOR to determine how patterns of drug-seeking overtime were altered by increased and eliminated arrestin-3 activity at the MOR.

## METHODS

### Mice

Mice of 3 genotypes were used in this study: 1) C57Bl/6 WT (n=20, 14 male, 6 female, 5 bred in-house and 15 purchased from the Jackson Laboratory) 2) RMOR (19) (n=15, 8 male, 7 female) bred in house, congenic >30 generations on C57Bl/6 and 3) Arr3-KO (15) (n=16, 7 male, 9 female) originally acquired from Dr. R. Lefkowitz (Duke University) (15) and bred in-house congenic for >30 generations on C57Bl/6. Adult mice aged 9-11 weeks at the start of training were used. Mice were singly housed with running wheels as extra enrichment upon entering the study and had access to food and water *ad libidum*. Single housing was necessary to monitor morphine consumption in the home cage. Mice were housed in a room with a reversed 12-hour dark/light cycle so that all study tasks took place during their active/dark period.

### Determination of physical dependence to oral morphine

Following exposure to orally available morphine (see figure 1A,B), mice were assessed for physical dependence to morphine. Mice were injected subcutaneously with 5mg/kg naloxone and observed in clear plexiglass chambers for signs of withdrawal including jumping, wet-dog shakes, teeth-chattering, and paw tremors. A global withdrawal score was calculated as the sum of these behaviors.

**Figure 1:**
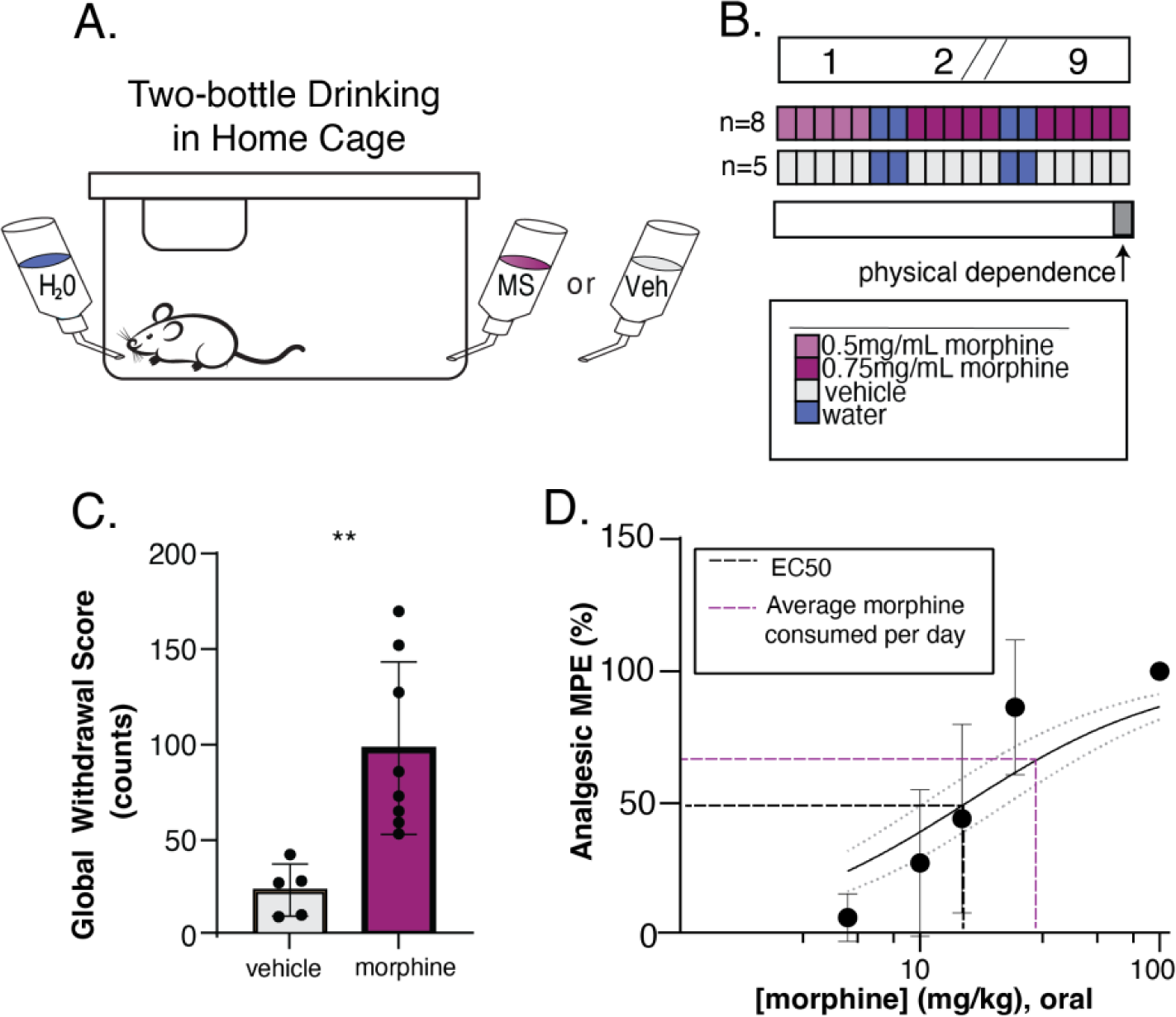
Oral consumption of morphine is sufficient to induce physical dependence and analgesia. A) Schematic of home cage setup with 24/7 water access and 24/5 morphine or 0.2% saccharin vehicle access. B) Experimental timeline to validate oral drinking exposure as a valid route of administration. Top bar shows time in weeks, where slashes indicate a repetition of previous weeks. Colored bars show available oral solutions in the home cage. Morphine at 0.5mg/mL (light purple) in 0.2% saccharin vehicle was available on the first week, then was increased to 0.75mg/mL (dark pink) in vehicle for the morphine drinking group (n=8). The vehicle solution alone (orange) was available to the saccharin drinking group (n=5). Mice had 24/7 access to water (blue), but two days a week the morphine or vehicle bottle was removed leaving the water bottle only. On the final day of exposure, naloxone precipitated withdrawal (physical dependence) was measured (gray bar). C) Physical dependence was assessed by injecting mice with 5mg/kg naloxone and calculating a Global Withdrawal Score for the subsequent 20-minute period (sum of jumps, wet dog shakes, teeth chatters, and paw tremors). The Global Withdrawal Score was significantly higher in morphine drinking mice as compared to their vehicle counterparts (p = 0.0047, two-tailed unpaired t-test). D) Analgesia was evaluated using a tail flick assay and a dose response curve was constructed to oral gavage of morphine (EC50 = 15.6) (black dotted line). The average amount of daily voluntary morphine consumption (31.3 mg/kg/day) (purple dotted line) is also visualized.

### Generation of oral dose-response curve to morphine

A dose-response curve to orally administrated morphine was determined using a radiant heat tail-flick assay (Tail-flick Analgesia Meter, Columbus Instruments. Columbus, OH). The light intensity was adjusted such that baseline latency (no drug present) to tail flick was 1.4-2.0 seconds. A maximum of three times the baseline latency (6.0 seconds) was used as a cutoff time to prevent tissue damage. A minimum of 5 independent subjects were tested for each dosing group. An oral gavage solution in sterile saline was prepared so that each subject received a maximum of 100µl when dosed by kilogram. Drug response latencies were measured 45 minutes following oral gavage of morphine solution. A non-linear fit equation in GraphPad Prism was used to determine the EC_50_ dose of oral morphine. Data are displayed as Analgesic Maximum Possible Effect (%MPE): 100*[(drug response latency−baseline latency)/(cutoff time− baseline latency)].

### Operant Training with saccharin reward

Med Associates operant conditioning chambers (Fairfax, VT) were used for the extent of this study. Mice were first trained to press a lever for a reward using saccharin as the reinforcer. Both active and inactive levers were present at the start of training. The active lever was indicated by the presence of a light cue above the lever while inactive levers were unlit. A press on the light-cued active lever delivered 15 µl of 0.2% saccharin sodium salt hydrate (Sigma-Aldrich, St. Louis MO) that was signaled by the illumination of a cue light above the delivery port and a 2.5-second tone (see Fig. 2A). Mice were trained in two stages: Stage 1 consisted of a progressive fixed ratio (FR) reinforcement schedule from FR1 (every active lever press produces a reward) to FR4 (four consecutive presses are required to produce a reward). Mice progressed to the next FR schedule after they obtained 20 rewards at each FR. To pass Stage 1 mice had to press a total of 200 times for 80 rewards (20 at FR1, 40 at FR2, 60 at FR3, and 80 at FR4). Each session lasted a maximum of 6 hours. Mice that failed to pass Stage 1 after 6 sessions were eliminated from the study. In Stage 2, mice were returned to the box for an FR1-FR4 progressive session with one reward at each FR step before progressing to the next step: admittance into the study. To pass Stage 2, mice had to press the active lever 10 times for 4 rewards (1 press at FR1, 2 at FR2, 3 at FR3, and 4 at FR4). Only mice that passed Stage 2 within one hour were entered into the study.

**Figure 2:**
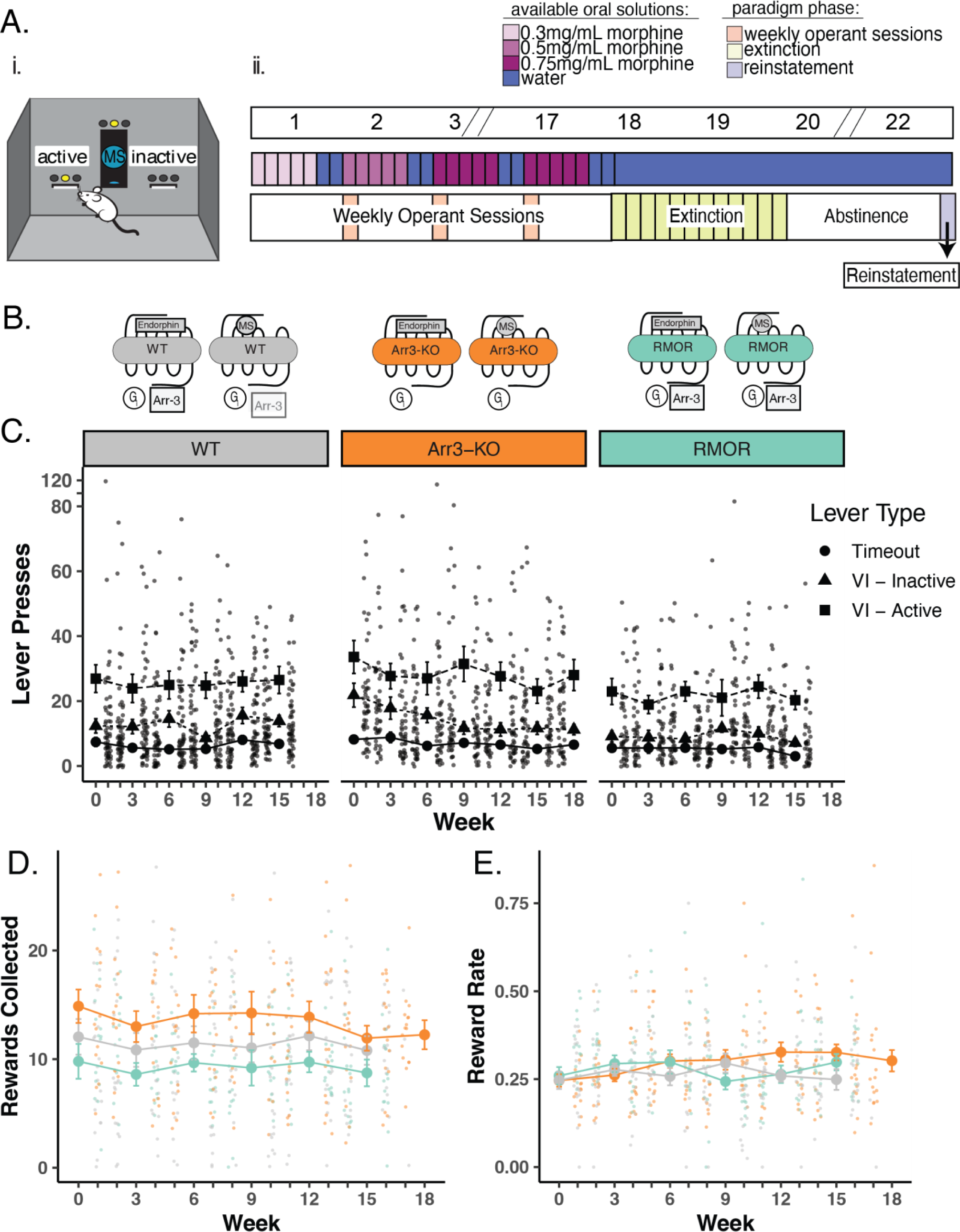
Deletion of arrestin-3 does not reduce drug-seeking behavior in an operant self-administration task. A) Experimental paradigm for longitudinal model of OUD. i. Schematic of operant self-administration chamber where lever pressing resulted in delivery or denial of a morphine reward. ii. Experimental timeline. Top bar shows example weeks where slashes indicate a repetition of previous weeks. Middle bar shows oral MS availability in the home cage where blue bars represent water alone and increasing concentrations of morphine (0.3 mg/mL, 0.5 mg/mL, and 0.75 mg/mL morphine) are lightest to darkest purple (middle bar). Mice were able to drink morphine *ad libitum* in their home cage (Fig. 1A) for five days a week and water seven days a week during the self-administration phase of the paradigm. Bottom bar shows the three phases of the paradigm. Phase 1: 16-19 weeks of home cage drinking, with an operant self-administration session (peach bars) one day per week. Phase 2: Lever pressing behavior was extinguished in up to 12 extinction sessions (green bars). Phase 3: Cue-induced reinstatement (light purple bar) of lever pressing following a 14-day period of complete morphine abstinence. B) Schematic of MOR signaling in WT (gray), Arr3-KO (orange), and RMOR (teal) mice in response to morphine and the endogenous ligand, endorphin. Effectors include G_i/o/z_ protein (G_i_, circle), Arrestin-3 (Arr3, square) C) Lever pressing behavior during operant self-administration phase in WT (gray), Arr3-KO (orange) and RMOR (teal) mice. Lever press counts are summarized (mean and standard error) for every three weeks of the self-administration phase with the distribution of individual subject counts displayed on the alternate weeks. Three types of lever press behaviors are described. Timeout (circles): any lever press that occurs in the first five minutes of a 30-minute session, Inactive (triangles): a press on an inactive lever during the final 25 minutes of a session, Active (squares): a press on an active lever during the final 25 minutes of a session. Only active lever presses could trigger reward delivery. D) Rewards collected during operant self-administration phase. Reward collection counts of WT, Arr3-KO, and RMOR (same colors as above) are summarized (mean and standard error) for every three weeks with the distribution of individual subject counts displayed on the alternate weeks. E) Reward rate during operant self-administration phase. Reward rate was calculated as rewards collected/total lever presses for each session. Session reward rates for WT, Arr3-KO, and RMOR (same colors as above) are summarized (mean and standard error) for every three weeks with the distribution of individual subject rates displayed on the alternate weeks.

### Oral Morphine Consumption Schedule

Mice who successfully completed operant training with saccharin were singly housed and their cages were outfitted with two bottles, one with water and the other with morphine sulfate (MS) (Mallinckrodt Pharmaceuticals, St. Louis, MO) + 0.2% saccharin to counteract the bitter taste of MS. In addition, to acclimate mice to the bitter taste of MS, the concentration of MS was 0.3 mg/mL in the first week and 0.5 mg/mL in the second week (Fig. 2Ai). After this, the concentration was increased to 0.75 mg/mL for the duration of the home cage drinking period. Mice had access to both the MS bottle and the water bottle 5 days per week and water only for the two days preceding each weekly operant session. MS and water bottles were weighed three times a week to monitor total morphine consumption.

### Operant Oral Self-Administration Weekly Schedule

After saccharin training was completed, mice remained on the same weekly schedule for 16-19 weeks (Fig 2B). After two days of access to only water, mice were placed in the operant box for a 30-minute session (peach bars, Fig 2Ai) that consisted of two distinct phases: a timeout period and a reinforcement period. The timeout period was signaled by the presence of a flashing light above the active lever and no light above the inactive lever. No lever presses were rewarded during this 5-minute timeout period, which in our OUD model reflects futile drug-seeking. After the 5-minute timeout, the light above the active lever stopped blinking and remained on, initiating the start of the 25-minute reinforcement period. During this period, the first active lever press was rewarded by delivery of a 15 µl oral morphine reward (0.5mg/mL MS in 0.2% saccharin), paired with the illumination of the light above the port and a 2.5-second tone. After that first reward, the wait time necessary between available rewards was unpredictable, from 1 to 90 seconds, but averaged 25 seconds. Time intervals for the variable interval reinforcement schedule were randomly selected from a 12-element Fleshler–Hoffman series to ensure all mice could access the same number of rewards (23). In our OUD model, a variable interval schedule was chosen to capture rates of lever pressing that reflect how hard a mouse is willing to work for drug since not all presses produce reward. All lever presses and all rewards consumed were automatically recorded during this weekly 30-minute session. After the operant self-administration session, mice were returned to their home cage with *ad libitum* access to both water and morphine for the next 5 days followed by two days of water access. This weekly schedule was repeated for 16-19 weeks.

### Extinction

Following 16-19 weeks of weekly operant self-administration, three 30-minute extinction sessions were conducted every day for a maximum of 12 days (Green bars, Fig 2B). Extinction sessions were identical to the self-administration sessions except that lever presses on the active lever never led to a morphine reward or the associated tone and light cues during any part of the session. Each mouse was assigned an individual extinction criterion delineated as an active lever press daily session average below 20% of their weekly session average during the final three weeks of their self-administration phase or four or fewer active lever presses, whichever number was higher. Once this criterion was met, the mouse moved on to the next phase of the paradigm. Mice moved on to the next phase (abstinence) after 12 days of extinction training regardless of lever pressing behavior. Some mice therefore had more extinction sessions than others. All lever presses during these extinction sessions were automatically recorded. During the extinction phase, mice had access to only water (no morphine) in their home cage.

### Abstinence and Reinstatement

Following extinction, mice were returned to their home cage with access to only water for two additional weeks with no morphine access (light purple bar, Fig 2B). Following this abstinence period mice were returned to the operant box for a single operant session. This session consisted of a 5-minute timeout period identical to previous sessions. After this timeout period, the light over the active lever remained on and a single non-contingent (no lever press required) morphine reward was delivered at the port with the associated light and sound cues. After this single non-contingent reward delivery, the light remained on over the active lever, but no additional rewards or cues were delivered. During this session, all lever presses, all head port entries, and the latency to collect the non-contingent reward were recorded.

### Calculation of compulsivity composite scores

Principle Coordinate Analysis and correlations analyses conducted in the R software packages factoextra (v 1.0.7) and corrplot (v 0.92) were used to identify measured behaviors through the paradigm indicative of drug abuse liability and that distinguish WT and RMOR mice from each other. A total of 16 measures from throughout the paradigm were selected to create a composite OUD/compulsivity score for each mouse (see Fig 4A for the variables used in the final score). The raw values for each mouse for each of these 16 measures were Z scored across the population of mice that completed the study (51 mice: 20 WT, 15 RMOR, 16 Arr3-KO). To give each phase equal weight when calculating the final score, a sub-score for each of the three phases (self-administration, extinction, reinstatement) was then created by averaging the Z scores of each behavioral measure in that phase for each mouse. The values for the operant self-administration phase came from the average of the final three weekly sessions for that mouse. The extinction values represented the average of the three sessions on each animal’s first day of extinction. A final compulsivity score was then created by adding the self-administration, extinction, and reinstatement sub-scores for each mouse. The distribution of composite compulsivity scores of WT mice were bimodal (R software package mclust), thus we used the mean and interquartile standard deviation (IQD) of WT compulsivity scores to determine categorical assignments of compulsive or non-compulsive for the entire population. The IQD is defined as the standard deviation of values between Q1 and Q3. All mice with a composite score of 1 IQD or more over the mean score of WT mice were designated as compulsive.

### Morphine Preference

On days 3-5 of the final week of the operant self-administration phase, we conducted a preference test for morphine (sweetened with 0.2% saccharin) versus saccharin alone. To do this, the water bottle in the home cage was replaced with a bottle of 0.2% saccharin for 4 hours during the dark cycle, and consumption of both morphine and saccharin was determined by weighing the bottles before and after this test. Preference for morphine over saccharin was calculated AS MS consumed (in mLs)/Total fluid consumed (in mLs).

### Statistical Analysis

All statistics were conducted using R and the RStudio software except for Figure 1B & D, which were constructed in GraphPad Prism software. Statistical tests were chosen based on the distribution of data in each group. Normality of data was assessed using a Shapiro-Wilk test and data with a p-value greater than 0.05 was considered normal. One-way ANOVA or t-tests were used to compare differences between groups where assumptions for normality and homogeneity of variance were met. The Kruskal-Wallis was used when assumptions of normality were not met.

### Study Approval

All protocols were approved by the Institutional Animal Care and Use Committee at the University of California Davis and are in accordance with the National Institutes of Health guidelines for the care and use of laboratory animals.

## RESULTS

### Oral morphine self-administration is sufficient to produce both analgesia and dependence

To emulate human-like patterns of OUD in a rodent model, we developed a paradigm that allowed mice to engage in naturalistic drug-taking with substantial drug exposure but also yielded sufficient information to quantify motivated drug-seeking behavior. To accomplish this, we utilized a combination of traditional operant self-administration and a variation of the two-bottle choice drinking task similar to a model we have described previously (22). To validate that the paradigm provides sufficient drug exposure, we examined whether voluntary drinking on this schedule was sufficient to produce opioid dependence in WT C57Bl/6 mice. Following 9 weeks of home cage morphine drinking (Fig. 1A,B), we evaluated mice for common effects of opioid withdrawal precipitated by naloxone injection (5 mg/kg). Mice that had access to morphine in their home cage had significantly higher global withdrawal scores than those who had access only to the 0.2% saccharine vehicle solution (p = 0.0047, two-tailed unpaired t-test) (Fig. 1C). In a separate set of mice, we also confirmed that oral morphine at doses comparable to daily voluntary morphine consumption was sufficient to produce analgesia in a tail flick assay (Fig. 1D).

### Deletion of arrestin-3 does not reduce drug-seeking behavior in a longitudinal OUD model

The WT MOR recruits arrestin-3 very weakly in response to morphine activation when compared with the recruitment promoted by endorphins/enkephalins (24) (Fig. 2B, gray). To determine whether the degree of arrestin-3 recruitment to the MOR modulates drug-seeking, we employed our longitudinal mouse model of OUD and used two have no ability to recruit arrestin-3 (Fig. 2B, orange). In RMOR mice, the receptor recruits arrestin-3 in response to both endorphin and morphine activation (Fig. 2B, teal).

To monitor the transition to compulsive drug-seeking and relapse as described previously (22), we implemented a paradigm which consisted of three separate stages: 1) Weekly Operant Self-administration 2) Extinction and 3) Reinstatement (Fig. 2Aii). Drug-seeking behavior was evaluated during each phase in an operant task (Fig. 2Ai) during which presses on an active lever may or may not yield an oral morphine reward on a variable interval reinforcement schedule. Mice were initially trained to press the lever for a saccharin reward and only mice who met the initial training criteria were advanced to the morphine-seeking task.

Mice in all three genotypes learned the task at equivalent rates and demonstrated a preference for the active lever over futile lever pressing (presses on an inactive lever or on any lever during the initial timeout period of the session). Lever pressing activity was stable through many weeks of self-administration sessions, and there were no significant differences from average WT lever pressing behavior in RMOR or Arr3-KO mice (determined by one-way ANOVA with Tukey’s multiple comparisons test) (Fig. 2C). On average, RMOR mice achieved fewer morphine rewards during operant sessions (Fig. 2D), but this was not statistically significant. When corrected for total lever pressing behavior, their reward rate was not different from the other two groups (Fig. 2E), reflecting that morphine is a more potent reinforcer in RMOR mice as previously reported (22).

Following the operant self-administration phase of the paradigm, mice were given extinction sessions three times daily in which cues and drug reward were no longer presented in response to active lever presses. Extinction sessions were ceased once a mouse met an individualized criteria determined as 20% of active lever pressing displayed during late operant sessions, or fewer than 4 lever presses in a session. Mice that reached 12 days of extinction training were automatically advanced to the next phase of the paradigm. The majority of mice extinguished their drug-seeking behavior within 12 days, but there was a significant effect of genotype on days to reach extinction (p = 0.036, F = 3.566, one-way ANOVA) as Arr3-KO mice took longer to reach extinction compared to the WT group (p = 0.028, Tukey’s multiple comparisons test) (Fig. 3A). 7 out of 16 (43.75%) Arr3-KO mice did not reach their extinction criteria within 12 days, something that only occurred in 2 (13%) RMOR and 1 (5%) WT mouse.

**Figure 3:**
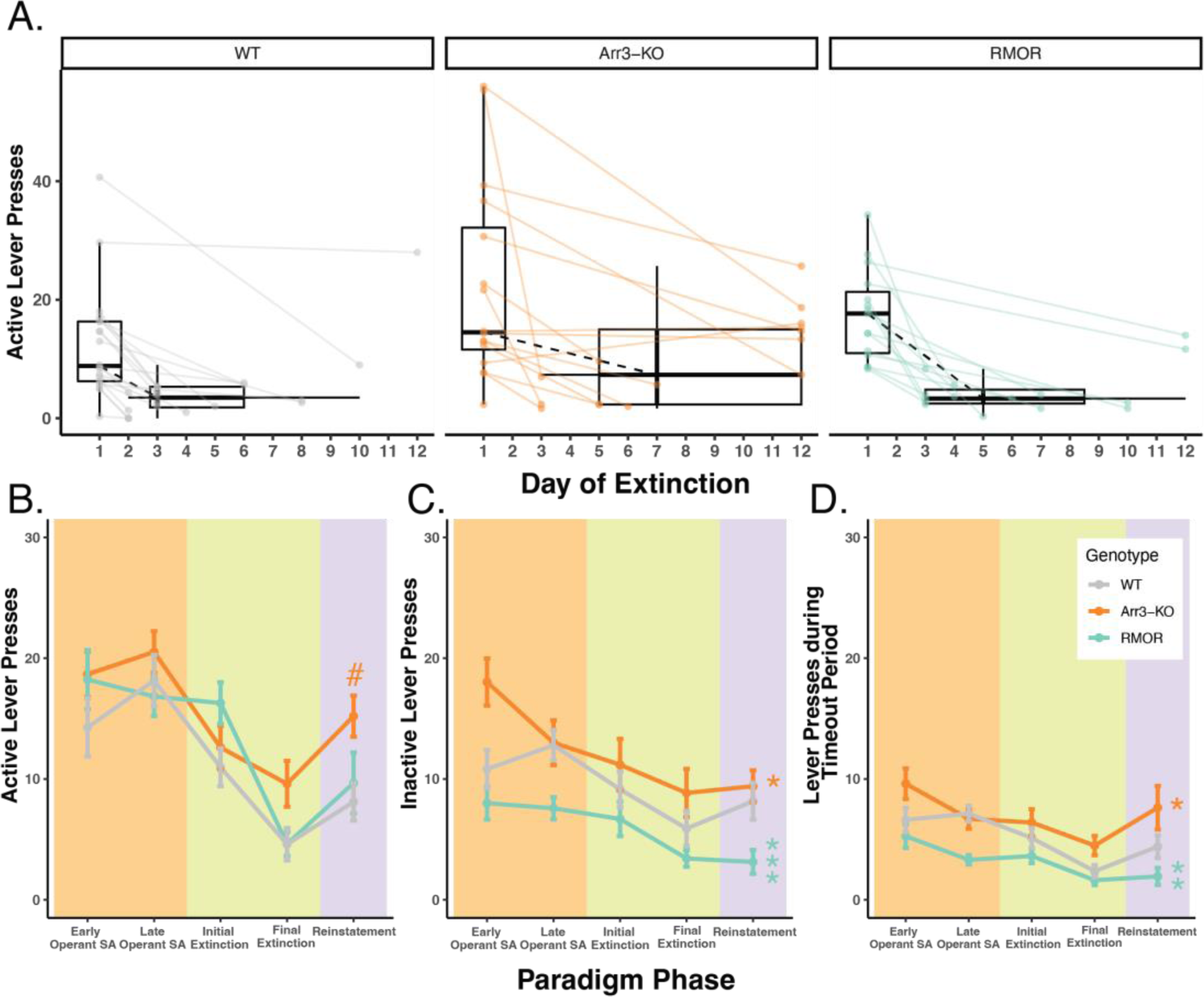
Deletion of arrestin-3 does not reduce drug-seeking behavior in a longitudinal model of OUD. A) Summary of active lever pressing during the extinction phase. Box plots and points represent the distribution of active lever press counts on the first day (Day 1) and final day (variable) of extinction in WT (gray), Arr3-KO (orange) and RMOR (teal) mice. Final day box plots also summarize the (horizontal) distribution of number of days to reach extinction which varied by mouse. Black dashed lines show the change in median lever press count between the first and median final day of extinction. Arr3-KO mice took significantly longer to reach extinction than WT mice (p = 0.028, One-way ANOVA with Tukey’s multiple comparisons test). B) Active lever presses during each paradigm phase. Each phase of the paradigm, self-administration (peach), extinction (green), and reinstatement (purple), is denoted by background colors. Within the self-administration phase the lever presses from the first three weeks (early) and final three weeks (late) are summarized separately. Within the extinction phase, the lever presses from the initial (Day 1) and final (variable) are summarized separately. Mean and SEM are shown for WT (gray), Arr3-KO (orange) and RMOR (teal) mice. Genotype significantly affected active lever pressing across the paradigm (p < 0.001, Kruskal-Wallis test) Arr3-KO mice showed significantly more active lever pressing than WT and RMOR during the reinstatement phase (p = 0.035 & p = 0.039 respectively, Dunn test). # indicates significant difference from WT within individual phase. C) Inactive lever presses during each paradigm phase. Data are displayed according to the specifications of B. Genotype significantly affected inactive lever pressing across the paradigm (p < 0.001, Kruskal-Wallis test). RMOR (p < 0.001) and Arr3-KO (p = 0.017) mice displayed significantly different lever pressing than WT (Dunn test). * indicates significant difference from WT of all data across phases. D) Lever presses during the timeout period for each paradigm phase. Data are displayed according to the specifications of B. Genotype significantly affected inactive lever pressing across the paradigm (p < 0.001, Kruskal-Wallis test). RMOR (p = 0.002) and Arr3-KO (p = 0.017) mice displayed significantly different lever pressing than WT (Dunn test). * indicates significant difference from WT of all data across phases.

After extinction, mice returned to their home cage for two weeks of abstinence with access to only water to drink. Following this abstinence period mice were returned to the operant box for a single operant reinstatement session. This session was identical to a single 30-minute extinction session except mice received a single non-contingent morphine reward at the termination of the timeout period. Genotype significantly affected drug-seeking behavior during reinstatement (p = 0.02, Kruskal-Wallis test) as Arr3-KO mice displayed more active lever pressing than WT and RMOR groups (p = 0.035 & p = 0.039 respectively, Dunn test) (Fig. 3B). This is likely because several Arr3-KO mice did not effectively extinguish their drug-seeking behavior. A Kruskal-Wallis test did not reveal a significant genotype effect in active lever pressing on the final extinction day (p = 0.068), but Arr3-KO mice pressed more than other groups reflecting their lack of extinction. Overall futile lever pressing (inactive lever pressing or lever pressing during the timeout period) was significantly affected by genotype (p < 0.001 for both futile lever types, Kruskal-Wallis test). RMOR mice displayed significantly less inactive (p < 0.001) and timeout (p = 0.002) lever pressing than WT despite no significant difference in their active lever pressing (p = 0.559, Dunn test) (Fig. 3C,D). Conversely, Arr3-KO mice had slightly more futile lever pressing behaviors overall (p = 0.017 for both futile lever types, Dunn test) compared to WT mice, though this is partially driven by their increased tendency to press the inactive lever early in the self-administration phase. Overall, these data show that while the RMOR phenotype may offer some protection from compulsive drug-seeking behaviors in this model, arrestin-3 deletion does not offer improved outcomes after prolonged morphine exposure and may increase compulsive drug-seeking.

### Arrestin-3 deletion does not improve compulsivity as defined by a behavioral composite score

OUD is a complex diagnosis that involves a combination of behaviors and varies in its individual presentation. Because our experimental paradigm was designed to encapsulate several addiction-relevant behaviors, we considered a multi-variate analysis strategy. A Principal Coordinate Analysis (PCoA) of 16 behavioral measures across our entire operant paradigm (Fig. 4A) revealed that RMOR mice clustered tightly, whereas both WT and Arr3-KO mice had high variability across both dimensions (Fig 4B). We posited that this variability could reflect a bifurcation of phenotype in the WT and Arr3-KO groups in which a subset of mice adopt a compulsive behavior pattern just as only a subset of humans exposed to opioids develop OUD.

**Figure 4:**
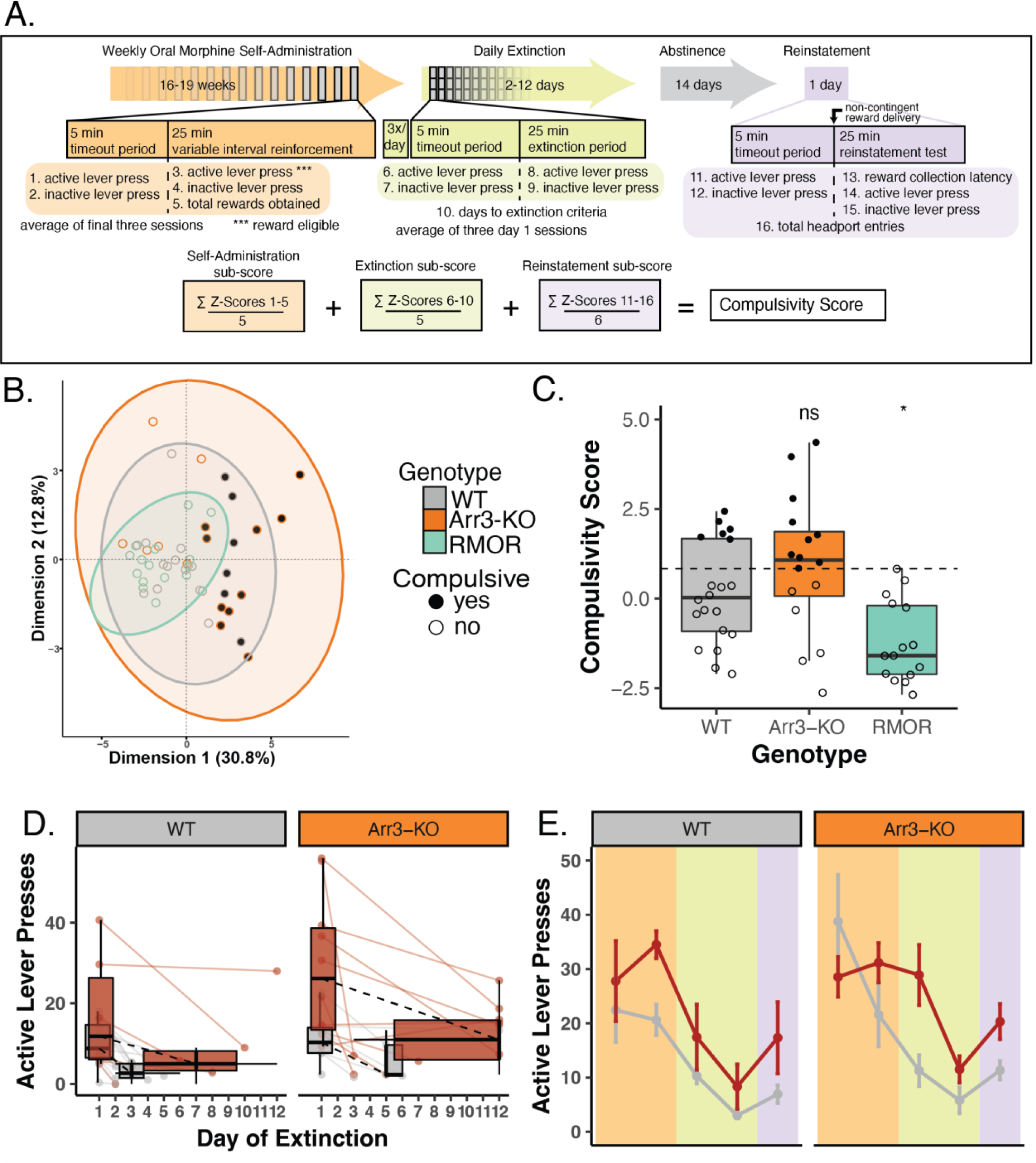
Arrestin-3 deletion does not improve compulsivity as defined by a behavioral composite score. A) Construction of composite behavioral score. Three paradigm phases, self-administration (peach), extinction (green), and reinstatement (purple, are displayed as a timeline with boxes below describing the details of a 30 minute task session. Variables included in the composite score are numbered 1-16 at their corresponding place in the paradigm timeline. The equation at the bottom of the panel displays the composite score calculation. B) Principal coordinate analysis was conducted using the 16 variables listed above. This revealed a tight cluster of RMOR (teal) mice while WT (gray) and Arr3-KO (orange) mice have more variable behavior. All but two of the compulsive mice (filled grey and orange circles) fall outside the RMOR cluster. C) Individual compulsivity scores for each mouse. Scores of compulsive mice (closed circles) were greater than one interquartile deviation above the mean composite score of WT mice. Non-compulsive mice (open circles) fell below this threshold. No RMOR mice were defined as compulsive. Scores of RMOR mice significantly differed from WT mice, but Arr3-KO mice did not in a one-way ANOVA with Tukey’s multiple comparisons correction (p = 0.031 and p = 0.262, respectively). D) Active lever pressing during the extinction phase in compulsive and non-compulsive mice. Individual subject data (points and solid lines) from figure 3A are revisualized with compulsive (red) and non-compulsive (gray) mice summarized (box plots) as distinct groups. Black dashed lines show the change in median lever press count between the first and median final day of extinction for each group. RMOR mice are not shown as they do not have a subset of compulsive animals. E) Active lever presses during each paradigm phase in compulsive and non-compulsive mice. Lever pressing summary data from figure 3B is revisualized with compulsive (red) and non-compulsive (gray) mice summarized (mean and SEM) as distinct groups. RMOR mice are not shown as they do not have a subset of compulsive animals.

We calculated a composite score for each mouse that incorporated all 16 measures used in the PCoA (Fig 4A). Mice were designated as compulsive if their composite score fell above a threshold determined as one interquartile deviation above the mean composite score of the WT group (Fig. 4C). By these criteria, of the 20 WT mice, 6 (30%) were compulsive. Of the 16 Arr3-KO mice, 10 (62.5%) were compulsive. None of the 15 RMOR mice were compulsive, replicating what we have previously shown (22). Comparison of composite compulsivity scores showed a significant genotype effect (Fig. 4C, p = 0.001, F = 8.007, one way ANOVA). In a Tukey’s multiple comparisons test, Arr3-KO mice had no significant difference in compulsivity score from the WT group (p = 0.262), but WT and RMOR mice show a significant difference in compulsivity (p = 0.031) (Fig. 4C). These data confirm our previous work indicating that effective arrestin-3 engagement diminishes the liability for compulsive drug-seeking. Further, they suggest that preventing arrestin-3 engagement does not reduce compulsive drug-seeking and it may even exacerbate it. This is a deviation from what we see with the physiological effects of analgesia and tolerance where both enhancement and elimination of the arrestin-3 pathway cause similar shifts. However, it aligns with the RMOR phenotype of reduced physiological and affective dependence that is absent in Arr3-KO mice.

When we re-visualize the lever pressing behaviors after separating the mice into compulsive and non-compulsive groups, we observe a divergence of activity that is not apparent when we examined genotype differences. Compulsive mice take several more days to extinguish their lever-pressing behavior (p = 0.012, Kruskal-Wallis test) (Fig 4D). They also show an apparent escalation in drug-seeking through the self-administration phase that is not echoed by the non-compulsive group. This is apparent in the divergence of active lever pressing which is significantly higher in compulsive mice at the end of the self-administration phase (p < 0.001) despite there being no difference between the same mice at the outset of the phase (p = 0.337, Kruskal-Wallis test) (Fig 4E). This is not surprising, as these variables are contained in or derived from those within the composite scores used to assign the compulsivity threshold. It does, however, affirm the hypothesis that drug-seeking phenotypes may be more appropriately treated as bimodal than just highly variable. This idea is bolstered by the tightly clustered variability of the RMOR mice, none of which were compulsive.

### Compulsive drug-seeking behavior is independent of morphine consumption or preference

The vast majority of the morphine consumption in our paradigm occurred during home cage drinking. Individual mice were highly variable in their weekly morphine consumption with a range of 2.09 to 11.1mgs consumed per week, on average. There was no significant difference in average morphine consumption (p = 0.799, Kruskal-Wallis test) between WT and Arr3-KO mice (Fig. 5A). There was no correlation in total morphine consumption and compulsivity score (p = 0.57, R = −0.097) (Fig 5B). There was also no significant difference in morphine consumption between compulsive and non-compulsive mice, (Fig. 5C, p = 0.239, Kruskal-Wallis test).

**Figure 5:**
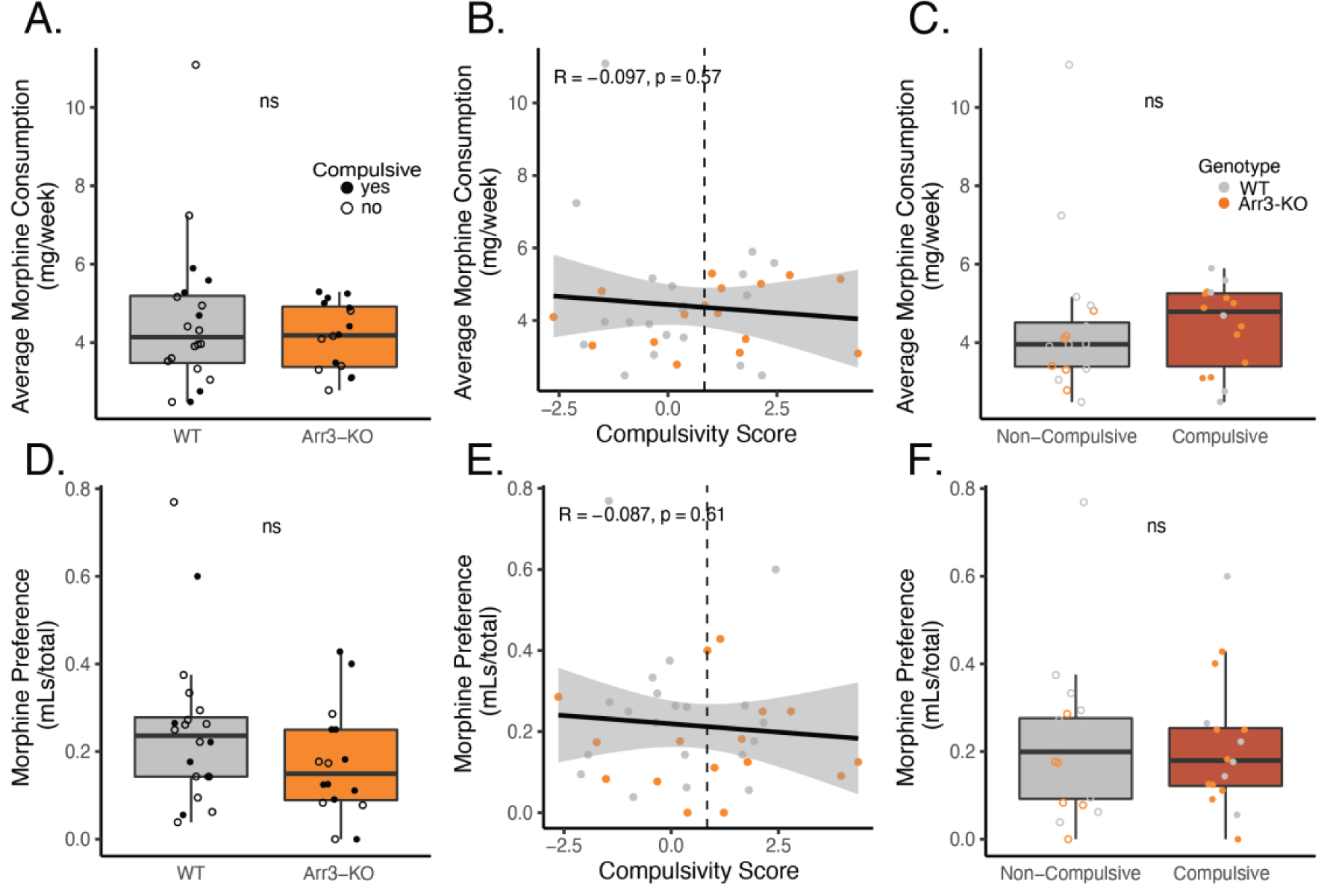
Morphine consumption or preference for morphine over saccharin does not predict compulsivity. A) Average weekly morphine consumption during the self-administration phase of the paradigm in WT (gray) in Arr3-KO (orange) mice. There is no significant difference between genotypes (p = 0.799, Kruskal-Wallis test). B) Average morphine consumption does not correlate with compulsivity score in a simple linear regression model (p = 0.57, R = −0.097). Vertical dashed line indicates compulsivity threshold score. C) Average morphine consumption does not differ between compulsive (red) and non-compulsive (gray) mice (p = 0.239, Kruskal-Wallis test). D) Preference for morphine over saccharin for WT (gray) and Arr3-KO (orange) mice. Preference for morphine in 0.2% saccharin vs 0.2% saccharin alone was measured on the final week of the self-administration paradigm in a 4-hour two-bottle choice test in the home cage. Preference (volume MS consumed/total volume consumed) did not vary significantly between genotypes (p = 0.156, Kruskal-Wallis test). E) Preference for morphine does not correlate with compulsivity score in a simple linear regression model (p = 0.61, R = −0.087). Vertical dashed line indicates compulsivity threshold score. F) Preference for morphine does not differ between compulsive and non-compulsive mice (p = 0.911, Kruskal-Wallis test).

Because individuals with OUD often display a preference for opioid drugs over other sources of positive affect, we also assessed whether drug-seeking behavior was related to voluntary consumption of morphine (a drug reward) over saccharine (a naturalistic reward). During week 17 of our self-administration phase of the paradigm, we measured morphine and saccharine consumption in a traditional two bottle choice test over a four-hour period. There was no significant difference in preference for morphine versus saccharine between WT and Arr3-KO mice (p = 0.156, Kruskal-Wallis test) (Fig. 5D). Preference for morphine did not correlate with compulsivity score (Fig. 5E, p = 0.61, R = −0.087) and there was no significant difference in morphine preference between compulsive and non-compulsive mice (p = 0.911, Kruskal-Wallis test). These data indicate that morphine consumption and preference alone are not predictive of liability for compulsive drug-seeking behavior. This aligns with the realities of human drug use in which many individuals engage in medical or recreational opioid use without developing OUD.

## DISCUSSION

### Arrestin-3-MOR activity does not cause or exacerbate compulsive drug-seeking

Here we use three genotypes of mice with different abilities to recruit arrestin-3 to the MOR to show that deletion of arrestin-3 does not protect against compulsive morphine-seeking in a mouse model of OUD. In a longitudinal paradigm that mimics human opioid consumption with a combination of *ad libitum* morphine access and contingent (motivated) drug-seeking, Arr3-KO mice displayed as much morphine-seeking behaviors as WT and RMOR mice. In addition to similar performance in the operant self-administration phase of the paradigm, WT and Arr3-KO mice consumed similar amounts of morphine and showed similar preference for morphine over naturalistic reward in non-contingent drug access contexts. When morphine reward was no longer available, Arr3-KO mice were slower to extinguish their drug-seeking behavior than the other groups. In fact, several Arr3-KO mice achieved the maximum number of extinction days and progressed through the paradigm without reaching their activity-based extinction criteria. This resistance to extinction may explain why the Arr3-KO mice had a stronger reinstatement effect. When we created a composite score to quantify compulsivity based on a multi-variate analysis of several behavioral outcomes, a subset of both WT and Arr3-KO mice were compulsive drug-seekers. In contrast, none of the RMOR knock-in mice exhibited drug-seeking behavior that reached the threshold for compulsivity (Fig. 4C).

These data make a clear case that loss of arrestin-3 activity does not protect against the behavioral components of OUD. While Arr3-KO mice were more vulnerable to developing some OUD-relevant behaviors in our paradigm, it is unclear whether this means that low arrestin-3 engagement by opioid drugs increases their abuse liability. The Arr3-KO is a global knockout, and these effects could be influenced by the arrestin-3 pathway at any number of other receptors. Although it is clear that deletion of arrestin-3 does not improve outcomes in an OUD model, it is possible that engagement of the arrestin-3 pathway offers some protection from abuse liability of these drugs as we have previously reported (22). No RMOR mice were classified as compulsive in this study nor do they develop analgesic tolerance to morphine under conditions where both WT (19, 20) and Arr3-KO (20) mice do. RMOR mice also do not show either physical (19) or affective (22) dependence whereas both Arr3-KO (16) and MOR 11S/T-A (18) mice show dependence at a similar or exacerbated level compared to WT. The development of tolerance and dependence presents major limitations to the clinical utility of opioids and are complimentary to the behavioral exemplars of abuse liability. These combined physiological and behavioral phenotypes in the RMOR mice justify a renewed interest in how arrestin-3 signaling might be exploited for opioid development strategies.

### Opioid reward is an insufficient indicator of abuse liability

In our paradigm, which spanned several months, neither morphine consumption nor morphine preference was predictive of compulsive drug-seeking. Motivation to seek drug as measured by self-administration behavior also did not determine compulsive drug-seeking. Our data overall indicate that compulsive drug-seeking is not driven by opioid reward alone. This suggests that many of the behavioral assays, including simple operant responding, conditioned place preference, and consumption traditionally used as addiction proxies may not be predictive of actual liability to misuse drugs. This is consistent with the observation that although morphine reward is enhanced in both RMOR (22) and Arr3-KO (21) mice compared to WT mice, RMOR mice do not transition to compulsive drug-seeking (22), while a subset of both Arr3-KO and WT mice do. As morphine is rewarding in all three of these genotypes (21, 22), these data indicate that future opioid drugs should be evaluated beyond their ability to produce reward with a more holistic understanding of abuse liability.

In humans, OUD is evaluated based on a diverse set of diagnostic criteria that encompass physiological, psychological, and social effects of opioid use (25). Though it is impossible to recapitulate all these criteria in an animal model, more care could be taken to appreciate the heterogeneity of the disease. We attempted this with a multi-variate method that employs a PCoA and considers behaviors measured in multiple phases of an extended OUD paradigm. This allowed us to categorize animals into compulsivity groups based on a calculation that assigns equal importance to several behaviors that may or may not ultimately be relevant to the individual. Many models of substance use and misuse are well-established in the field, all of which have a role in unraveling the mechanisms of substance use disorders. Given the complexity of these disorders, it is in the interest of the field to adopt analytical approaches capable of simultaneously considering multiple animal behavioral outputs and how they may interact. We give one example here, but other techniques such as machine learning or meta-analyses would also be useful in evaluating these complex phenotypes.

### Balanced agonism is an under-studied strategy with potential to improve opioid therapeutics

The differentiating characteristic of RMOR mice is that MOR signaling has been altered to reflect that of the endogenous opioids, as the MOR recruits arrestin-3 and is endocytosed and recycled in response to morphine (26). This is not the case with the WT MOR which only recruits arrestin-3 when GRKs or arrestins are overexpressed (6, 11). In neurons, opioid peptides, but not morphine, promote MOR endocytosis (13, 27), a consequence of arrestin recruitment. Likewise, *in vivo*, morphine administration also produces little endocytosis (26, 28) compared to opioid peptides (29–31). Our data from RMOR and Arr3-KO mice imply that G protein-biased opioid ligands which do not engage arrestin-3 will not prevent abuse liability. This is a critical finding as opioid drug development has focused on the development of ultra-G-biased ligands for the past two decades. TRV-130 (Oliceridine) is one of these ligands and was FDA-approved in 2020—the first new opioid in 4 decades. This push to develop ultra-G-biased ligands followed reports that Arr3-KO mice show increased analgesia (15) and reduced tolerance (16) and respiratory depression (17) in response to morphine compared to WT mice, indicating that biased ligands could ameliorate these key side effects. However, several recent reports have failed to reproduce these findings in Arr3-KO mice (20, 32) and clinically, Oliceridine did not significantly reduce respiratory depression (33). As no studies have assessed the abuse liability of the new ultra-biased ligands, our results indicate clinically relevant risks that should not be ignored. These findings, coupled with the recent reports on respiratory depression, indicate that ultra G-biased ligands are unlikely to improve on existing side effect risks. For all these reasons, we posit that more work is needed to assess the benefits of a signaling profile that resembles endogenous opioids (2).

We describe endogenous opioid signaling as balanced because it effectively engages both the G protein and arrestin-3 pathways. Recapitulating balanced signaling with exogenous ligands is immensely challenging. Categorizing ligands as balanced or biased depends on quantification of arrestin-3 recruitment and signaling to G_i/o/z_ G protein effectors, techniques which are highly disputed and rife with caveats. This has made it difficult to assign a single signaling bias value for morphine—though it is always more G-biased than the endogenous ligands regardless of GRK/arrestin levels (34–36).

The only clinically-utilized opioid drug that approaches a signaling balance comparable to endorphins and enkephalins is methadone (28). No other existing balanced tool compounds have been tested *in vivo* because they have low potency (37), poor solubility (38), or were abandoned in favor of ultra G-biased ligands. In preclinical models, methadone produces less tolerance and less dependence than morphine (28). Though it is rarely used as a first line analgesic in humans because of its highly variable half-life, a few controlled studies show reduced tolerance to methadone in humans (see review for studies within) (39). However, methadone differs from morphine not only in pharmacokinetics but in many aspects of pharmacology (40). This makes it difficult to say that bias is the primary factor in its reduced side-effect profile. It would be informative to examine methadone tolerance, dependence, and compulsive drug-seeking in a mouse model that cannot recruit arrestin-3 to the MOR, such as the MOR 11S/T-A knock-in mouse (18). This effective conversion of methadone into a biased agonist would complement the findings from the RMOR mouse where morphine performs as a balanced agonist.

The phenomenon of the opioid epidemic demands multiple angles of attack. The phenotype of the RMOR mice gives hope that opioid agonists which provide both analgesia and reward without precipitating OUD could still be attainable. This goal remains vital as no alternative drugs exist for the treatment of severe pain. It is past time to expand our strategies in the areas of pharmacology and drug development and to meet this challenge with a tenacity that rivals that of this public health crisis.

## ACKNOWLEDGEMENTS

The authors would like to thank Dr. Robert Lefkowitz for the arrestin-3 knock out mice and all members of the J. Whistler and C. Whistler labs for their valuable input on the manuscript and their ongoing support. We would also like to thank Dr. Rishidev Chaudhuri for his thoughtful evaluation of our manuscript.

## AUTHOR CONTRIBUTIONS

JLW, LF and SWG designed the experiments. LF, ZR, SWG, JG, AG, MK, and ND performed experiments. Data analysis and interpretation was performed by SWG, RF, LF, CW, JLW, KI, and IS. SWG, JLW, LF, RF, and IS designed and generated the figures. SWG, JLW and LF wrote the manuscript. All authors contributed to manuscript editing.

## FUNDING AND COMPETING INTERESTS

This study was funded by the National Institutes of Drug Abuse under award numbers F31DA056222 (LF) and R21DA049565 (JLW, CAW), R01DA055708 (JLW), F31DA051116 (SWG), and by funds provided by the state of California as start up to JLW. The authors declare no competing interest.

## Notes

### Competing Interest Statement

The authors have declared no competing interest.

### Summary of Updates

We discovered mistakes in the original analysis pipeline that have been corrected here. We have also added additional figures and analyses to the manuscript that were not in the original version. Author order has changed to reflect the contributions for this revised version.

